# Diversity of translation initiation mechanisms across bacterial species is driven by environmental conditions and growth demands

**DOI:** 10.1101/167429

**Authors:** Adam J. Hockenberry, Aaron J. Stern, Luís A.N. Amaral, Michael C. Jewett

## Abstract

The Shine-Dalgarno (SD) sequence is often found upstream of protein coding genes across the bacterial kingdom, where it enhances start codon recognition via hybridization to the anti-SD (aSD) sequence on the small ribosomal subunit. Despite widespread conservation of the aSD sequence, the proportion of SD-led genes within a genome varies widely across species, and the evolutionary pressures shaping this variation remain largely unknown. Here, we conduct a phylogenetically-informed analysis and show that species capable of rapid growth have a significantly higher proportion of SD-led genes in their genome, suggesting a role for SD sequences in meeting the protein production demands of rapidly growing species. Further, we show that utilization of the SD sequence mechanism co-varies with: i) genomic traits that are indicative of efficient translation, and ii) optimal growth temperatures. In contrast to prior surveys, our results demonstrate that variation in translation initiation mechanisms across genomes is largely predictable, and that SD sequence utilization is part of a larger suite of translation-associated traits whose diversity is driven by the differential growth strategies of individual species.

## Introduction

Translation of a given messenger-RNA (mRNA) into functional protein relies on the ability of the translational apparatus to recognize the proper start codon. Bacteria have evolved several distinct mechanisms to discriminate between potential start codons, with the Shine-Dalgarno mechanism being the most ubiquitous [1, 2]. Variants of the Shine-Dalgarno (SD) sequence are frequently found upstream of bacterial start codons and functions to facilitate translation initiation by hybridizing with the complementary anti-SD (aSD) sequence on the 16S rRNA of the small ribosomal subunit (Fig. 1A).

**Figure 1:**
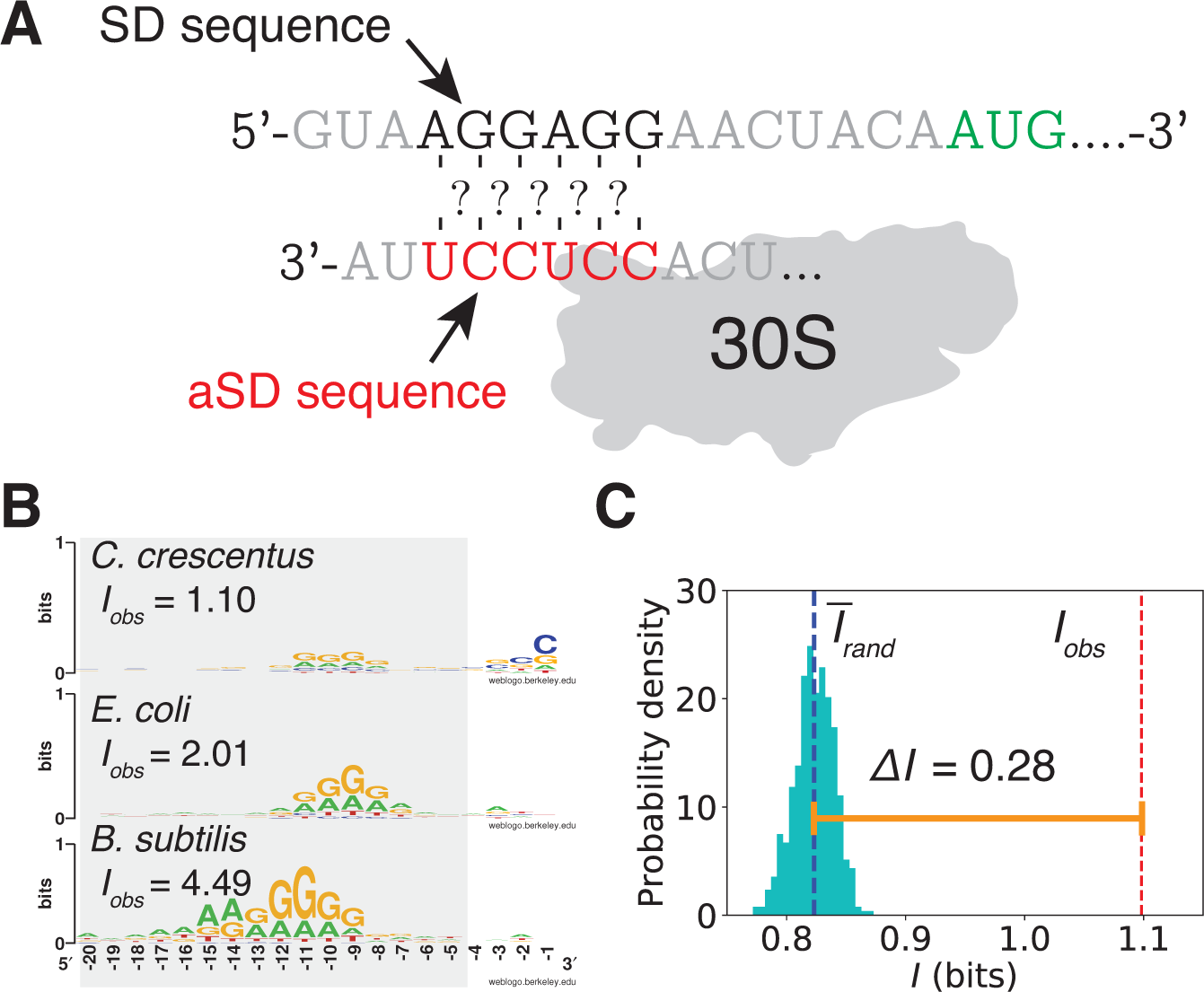
Sequence entropy quantifies genome-wide SD sequence utilization. (**A**), Illustration of the anti-Shine-Dalgarno(aSD)::Shine-Dalgarno(SD) sequence mechanism of translation initiation. (**B**), Representative sequence logos for three species derived from aligning the 5′ upstream region of all annotated coding sequences for individual genomes displays heterogeneity in sequence entropy. (**C**), Illustration of the Δ*I* metric for *C. crescentus*.

Variations of the canonical SD sequence occur across nearly the entire bacterial kingdom, and the aSD sequence is highly conserved (though notable exceptions exist) [3–9]. The importance of the SD sequence is further supported by the fact that SD-like sequence motifs are under-represented within the coding sequences of most bacteria, possibly reflecting their role in translational pausing and/or erroneous initiation [10, 11]. Like the diversity of SD sequence utilization, the degree of this under-representation is highly variable across bacterial species [12, 13].

For a given gene within an organism, it is known that the structural accessibility of the SD sequence, the thermodynamic binding potential between the SD sequence and the aSD sequence, and the exact positioning of the SD sequence relative to the start codon, are all features that collectively modulate the translation initiation rate of downstream genes [14–24]. Despite an abundance of research showing that the SD sequence enhances translation initiation and start codon recognition of downstream genes, there are several SD sequence-independent mechanisms that operate in bacteria including leaderless translation and RPS1-mediated translation of unstructured mRNA sequences [25–35]. Further, recent research suggests that mechanisms traditionally associated with eukaryotic species such as translational scanning and internal ribosome entry sites may also operate in bacterial systems [36, 37].

Given the high conservation of the aSD sequence—the reason for such diversity in translational mechanism utilization across species is puzzling [18, 19, 23]. For example, roughly 90% of *Bacillus subtilis* genes are preceded by a SD sequence while for *Caulobacter crescentus* the comparable number is closer to 50% [2, 5, 38]. Cross-species variation in translation initiation mechanisms may impact genetic isolation and transfer of genetic material, and quantifying the source and extent of variation may prove useful in identifying important genes in a genome or microbial community [9,39]. Further, the synthetic biology community is increasingly targeting both translation-system engineering and biotechnology applications involving less well-studied microbial species [40–45]. A better understanding of the factors shaping the utilization of different translation initiation mechanisms will aid in the design of synthetic gene constructs.

Here, we conduct a phylogenetic comparative analysis in order to isolate independent evolutionary events and show that the proportion of SD-led genes within a genome is strongly related to the growth demands faced by individual species. We develop a metric grounded on sequence entropy that captures the presence of over-represented motifs in the UTRs from a given genome, and demonstrate a link with protein production demands by showing that this metric is predictive of minimum doubling times for 187 bacterial species. Furthermore, we assemble a database of 613 phylogenetically diverse bacterial species and show that genome-wide variation in SD sequence utilization co-varies along-side a number of genomic features previously indicated to serve as markers for the translational burden imposed by rapid growth.

## Results

### Sequence entropy and its relation to SD sequence utilization

Several techniques have been previously used to quantify the overall utilization of the aSD::SD mechanism within a given species. In motif-based methods, researchers predefine several sub-sequences closely related to the canonical SD sequence and search a sequence window upstream of each protein coding genes within a given genome to determine the fraction of genes that are preceded by a SD motif [9,29]. Similarly, in aSD sequence complementarity based methods, researchers predefine a range upstream of the start codon to consider for each gene, a putative aSD sequence, and a hybridization energy threshold for determining whether a gene is SD-led or not [2, 4, 5, 7, 8].

Both of these methods rely on critical assumptions that may not hold when applied across large sets of phylogenetically diverse organisms. First, both of these methods carry an assumption that a SD sequence, regardless of its location relative to the start codon, has the same impact on translation initiation. However, experimental approaches have shown that spacing between the SD sequence and start codon can have dramatic effects on translation initiation rates [14–16, 18]. Second, both methods rest on a dichotomy between SD-led and non-SD-led genes. While this simplification is useful for *describing* the phenomenon, an abundance of research has shown that there are not two distinct categories but rather a spectrum of sequence complementarity that affects translation initiation in a continuous manner [16,18]. Third, bacterial genomes span a range of GC contents, and previous research has shown that it is critical to compare the proportion of SD-led genes in a genome to an appropriate null model expectation [2]. We therefore define the following term to summarize SD sequence utilization using the SD-motif based method by:

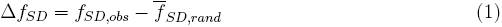

where *f*_*SD*_ is the fraction of genes within a genome classified as SD-led and *f*̅_*SD,rand*_ is the expected fraction of SD-led genes derived from repeating this calculation for 500 nucleotide shuffled ‘genomes’ and taking the average of these values (see Materials and Methods). We similarly define Δ*f*_*aSD*<−4.5_ to denote the same calculation as above, where SD-led genes are defined via hybridization of the putative aSD sequence using a threshold binding energy value of −4.5 kcal/mol (see Materials and Methods).

Additionally, we sought a complementary approach that would allow us to investigate hundreds of diverse genomes without having to *a priori* define an aSD sequence or SD motifs. For each genome we extract the 5′ upstream sequences from all annotated protein coding sequences (see Materials and Methods). We then sum the information contained in the sequences for this set within the region where SD-type motifs are expected to occur (−20 to −4 relative to the start codon):

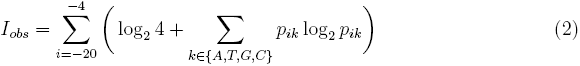

where *p_ik_* is the probability of finding base *k* at position *i*. We repeat this process for 500 shuffled genomes and compare the sequence information from the actual genome to the average of the nucleotide shuffled ‘genomes’:

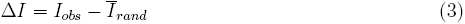

By definition, Δ*I* is a measure of non-randomness in the translation initiation region for a particular genome, which requires a single assumption: the existence of a predefined range upstream of start codons to include in the analysis. Eqs. (2) and (3) are agnostic to *which* sequence motifs are over-represented—thus alleviating the need to predefine putative aSD or SD motifs, which is necessary for the other methods discussed above. Figure 1B displays sequence logos of 5′ UTRs for three species to highlight the variation across species and Figure 1C illustrates our approach graphically.

We compiled a dataset of 613 bacterial species, unique at the genus level, for which we have complete, annotated genome-sequences as well as a high-quality phylogenetic tree describing their relatedness [46] (see Materials and Methods). In Figure 2A we show that while summary methods based on SD-motif and aSD sequence complementarity (Δ*f*_*SD*_ and Δ*f*_*aSD*<−4.5_, respectively) are linearly related for a large set of diverse species, there is a change in the slope that occurs for the Firmicutes phylum. Δ*I* also correlates strongly with these methods (Fig. 2A, Supplementary Fig. S1). However, in the Bacteroidetes phylum, we observe significant variation in Δ*I* without any apparent variation in either of the other metrics. These findings are consistent with prior research that identified changes in the aSD sequence region of the 16S rRNA sequence within this phylum [3]. The fact that the Δ*I* metric quantifies utilization of the aSD::SD mechanism for Bacteroidetes allows us to incorporate them into future analyses (Fig. 2A, red data points). Consistent with prior research [2], we show that SD sequence utilization according to the Δ*I* metric varies considerably across species while showing strong phylogenetic patterns (Fig. 2B).

**Figure 2:**
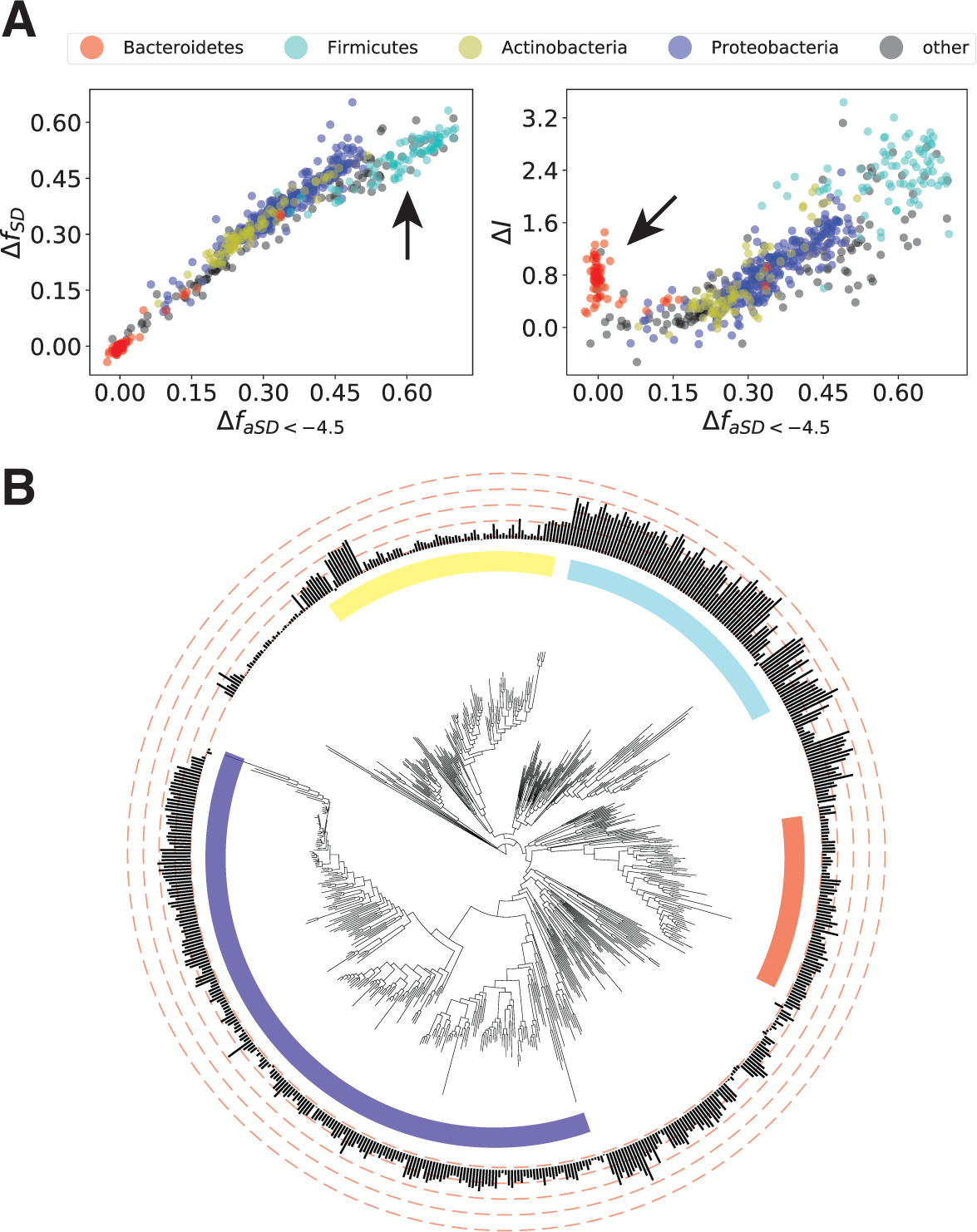
Relationship between Δ*I* and existing metrics of SD sequence utilization. (**A**), Comparison between different ways of summarizing SD sequence utilization, each data point represents a single genome. On the left, we show the relationship between SD motif and aSD sequence complementarity based methods (Δ*f*_*SD*_ and Δ*f*_*aSD*<−4.5_). On the right, we compare Δ*I* and Δ*f*_*aSD*<−4.5_. The four largest phyla are color-coded according to the legend. Arrows highlight phyla with ‘anomalous’ patterns. (**B**), Phylogenetic tree illustrating variation in SD sequence utilization across species according to the Δ*I* metric. Note the strong similarity in Δ*I* values for closely related species.

### Translation initiation and organismal growth demands

In previous research, Vieira-Silva *et al.* (2010) curated a list of minimum doubling times from the literature for a large number of bacterial species [47, 48]. Organisms that are capable of rapid growth have high protein production demands during these periods and there are a number of regulatory points that can be bottlenecks for this process. Meeting high translational demands associated with rapid growth requires coordination of a number of processes, and Vieira-Silva *et al.* (2010) showed that increasing numbers of rRNA and tRNA genes, and increasing codon usage biases amongst mRNAs in individual genomes were all partially predictive of the minimum doubling times of individual species.

At the individual gene level, translation initiation is an important control point, and we reasoned that translation initiation related features may similarly play an important role in meeting protein production demands imposed by rapid growth rates. We thus investigated whether variation in SD sequence utilization or the percentage of ‘ATG’ start codons were similarly predictive of minimum doubling times. We first replicate several of the findings of Vieira-Silva *et al.* (2010) using Phylogenetically Generalized Least Squares regression [49] to account for the lack of independence in species (see Materials and Methods). We verified that rRNA gene counts, tRNA gene counts, and a measurement of relative codon usage bias (a method based off the ‘effective number of codons’ (Δ*ENC*), see Materials and Methods) all have a highly significant relationship with minimum doubling times after controlling for phylogenetic effects (F-test, *p* < 0.002 for all cases, Table 1).

**Table 1:** Contribution of several factors for predicting minimum doubling times. The left column indicates individual variables that we considered for predicting minimum doubling times with the full multi-variate model listed at the top. *R*^2^ illustrates the overall goodness-of-fit for individual factors of minimum doubling time (*** indicates *p* < 0.001, ** indicates *p* < 0.01). Pagel’s λ is the fitted phylogenetic signal parameter, which we show with 95% confidence intervals in brackets. Values of λ close to ‘1’ indicate a strong phylogenetic signal in the residuals whereas a value close to zero indicates that there is no phylogenetic signal present in the residuals. The right column illustrates the change in goodness-of-fit from a model that includes all predictors to one that excludes only the variable of interest. Bold numbers in this column indicate variables with significant coefficients in the full multi-variate model (*p* < 10^−5^).

| Model for<br>min. doubling time | $R^2$ | Pagel's $\lambda$ | $\vert \Delta R^2\vert $ |
| --- | --- | --- | --- |
| Full model | 0.31*** | 0.93<br>[0.83,0.97] | - |
| $\Delta ENC$ | 0.17*** | 0.96<br>[0.92,0.99] | **0.16** |
| $\Delta I$ | 0.11*** | 0.97<br>[0.93,0.99] | **0.11** |
| 16S gene counts | 0.06** | 0.98<br>[0.95,0.99] | 0.02 |
| tRNA gene counts | 0.06*** | 0.98<br>[0.95,0.99] | 0.01 |
| ATG start % | 0.02 | 0.98<br>[0.95,0.99] | < 0.01 |

Next, we turn to translation initiation related metrics. We find that Δ*I* significantly correlates with minimum doubling times in this set of species (*p* < 10^−5^), showing the 2nd strongest correlation of any individual trait that we considered (Table 1). In contrast, we find that the proportion of protein coding genes containing an ATG start is not significantly correlated with minimum doubling times (*p* = 0.056).

In order to test the robustness of these findings and to assess overall predictability of minimum doubling times, we construct a multi-variable Phylogenetic Generalized Least Squares regression model that combines all of the listed factors, and find that only SD sequence utilization (Δ*I*) and relative codon usage biases (Δ*ENC*) have statistically significant coefficients (*p* < 0.001, both cases). Overall, a model containing all factors resulted in *R*^2^ = 0.31 (*p* < 10^−12^, Supplementary Fig. S2), while a more parsimonious model containing only the two factors with statistically significant coefficients resulted in *R*^2^ = 0.29 (*p* < 10^−13^). Removing either codon usage biases or Shine-Dalgarno sequence utilization from the full model substantially reduces its predictive power as illustrated in the right column of Table 1. In order to compare our work with prior research, we also conduct a phylogenetically *agnostic* linear regression model using all of these factors, which yields *R*^2^ = 0.57 (*p* < 10^−15^)—though we caution that ignoring the effects of shared ancestry will substantially bias statistical analyses, generally leading to inflated correlations and a high false positive rate. We also generated the same data as in Table 1 using Δ*f*_*aSD*<−4.5_ as a metric of SD sequence utilization and found largely similar results with less predictive power overall (Supplementary Table S1).

### Relationship between SD sequence utilization and other translation efficiency associated traits

Since a coordinated effort between multiple translational processes is required to maximize protein production, we reasoned that the various traits associated with efficient translation are likely to co-vary across species. In order to test this hypothesis, we assess the correlation between different definitions of SD sequence utilization and all of the alternative traits listed in Table 1 via Phylogenetic Generalized Least Squares regression. In Figure 3A we show the results of this analysis, finding that in all cases where a pair of traits is significantly correlated, the correlation is positive. Increasing SD sequence utilization is thus significantly associated with an increasing fraction of ATG start codons, increased 16S rRNA gene counts, increasing codon usage bias in ribosomal proteins, and increasing tRNA gene counts.

**Figure 3:**
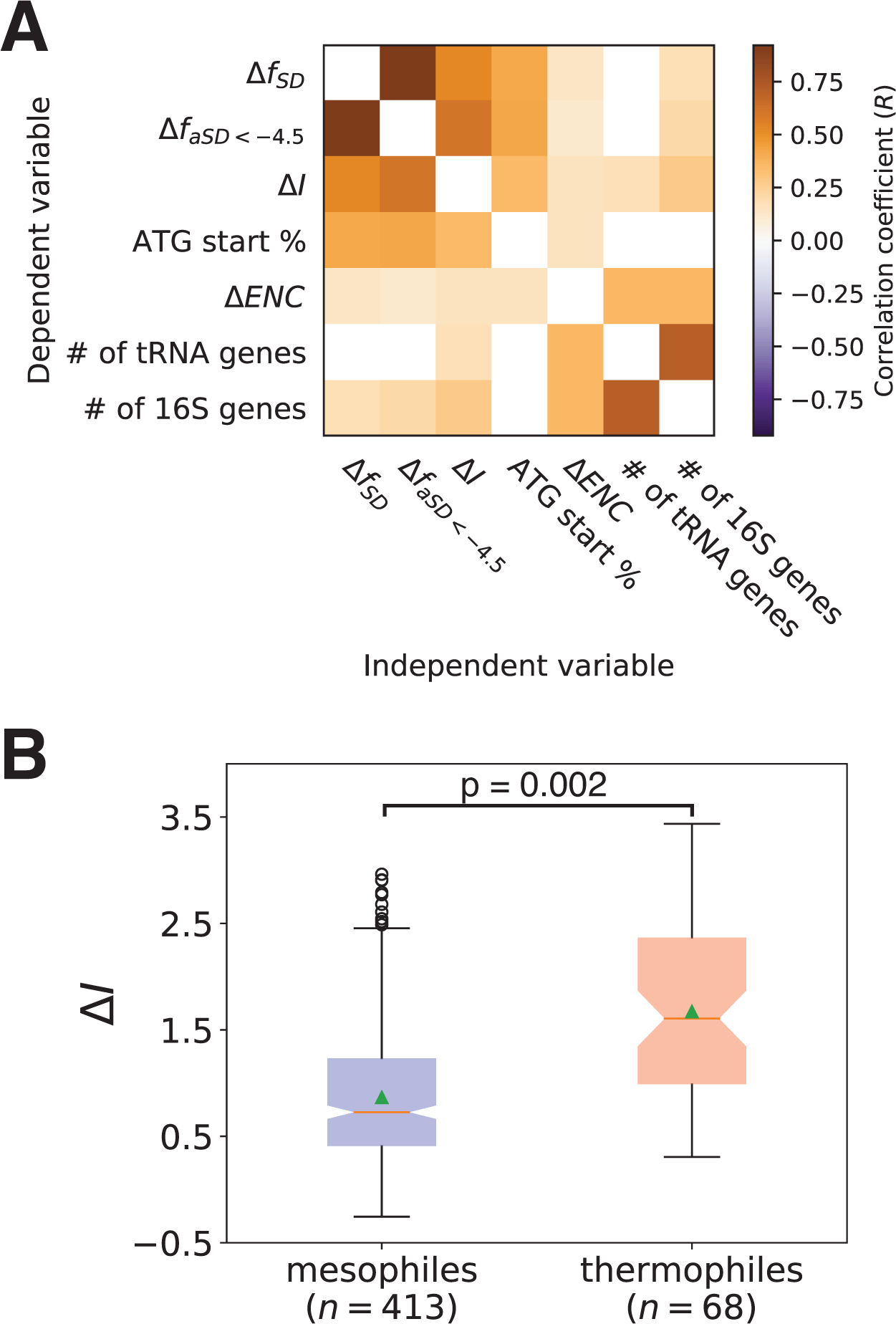
SD sequence utilization co-varies alongside a suite of translation-related traits and according to optimal growth temperatures. (**A**) Correlation matrix between listed variables used in Table 1 for a set of 613 diverse bacterial species shows that all features co-vary with one another in the positive direction for all significant cases. (**B**) SD sequence utilization, quantified using Δ*I* is significantly higher in thermophiles than in mesophiles.

We next test the overall robustness and universality of these results by independently analyzing these relationships within individual phyla. We specifically look at the 4 largest phyla in this dataset—Proteobacteria, Firmicutes, Actinobacteria and Bacteroidetes—and repeat the analysis from Fig. 3A. Again, we observe that every significant correlation that we observe is in the positive direction (Fig. S3). Notably, this phyla level analysis also highlights the advantage of the Δ*I* metric. When looking at relationships between variable SD sequence utilization in the Bacteroidetes phylum, Δ*I* shows significant relationships with three of the four other variables whereas Δ*f*_*SD*_ and Δ*f*_*aSD*<−4.5_ show no significant relationships apart from with one-another.

### Relationship between translation initiation mechanisms and optimal growth temperature

Having established that genome-scale SD sequence utilization is part of a suite of traits related to differential organismal growth strategies, we last wanted to assess whether other ecological factors relating to an organisms habitat may play a role in further constraining the evolutionary pressures related to SD sequence utilization. Specifically, we reasoned that since the aSD::SD sequence mechanism operates via RNA base-pairing, stronger sequence pairing would be necessary in order to get an equivalent level of stabilization of the translation initiation complex at higher temperatures.

Nakagawa *et al.* (2010) investigated this possibility, but found no association between SD sequence utilization and optimum growth temperatures [2]. By contrast, our phylogenetically informed modeling approach applied to this larger dataset (481 of the 613 species in our dataset have high-confidence growth temperature annotations) finds that temperature constrains genome-wide SD sequence utilization. Specifically, the genomes of thermophilic species display significantly larger values of Δ*I* than mesophilic species (Fig. 3B, F-test *p* = 0.002 using temperature as a fixed-effect in Phylogenetically Generalized Least Squares modeling). This finding further illustrates the role that ecological factors relating to growth conditions places on the evolution of genome architectures.

## Discussion

We have shown that variation in bacterial translation initiation mechanisms are a result of the differential growth strategies and environmental demands faced by individual species. We found that minimum observed doubling times and SD sequence utilization at the genome-scale are significantly correlated (*R*^2^ = 0.11). In a diverse dataset of 613 species, we further showed that SD sequence utilization predictably co-varies with several other genomic and environmental features, including the number of rRNA genes and optimal growth temperatures. Taken together, our findings demonstrate that organisms with greater translational demands are likely to co-evolve a common suite of genomic features that help to maximize translation during periods of rapid growth, and that SD sequence utilization is an important component of this shared genome architecture.

Our analysis throughout is performed in a manner that corrects for the confounding effects of shared ancestry between species, and our phyla specific results illustrate several critical points. First, the sign on the relationships between features that we observe is extremely robust, regardless of the phylum or SD sequence utilization summary statistic under consideration (Fig. 3, Supplementary Fig. S3). Increasing 16S gene counts, codon usage biases in ribosomal protein genes, tRNA gene counts, and ATG start codon usage fraction are universally associated with increasing SD sequence utilization. Second, Δ*I* is measuring an aspect of translation initiation region sequence preferences in the Bacteroidetes phylum that is not captured by previous models, which likely reflects novel sequence preferences in this lineage, a finding in need of further investigation. Future research with larger datasets may allow researchers to uncover branches within phylogenetic trees where mechanistic differences in the translational apparatus—resulting in differences in the slope and/or sign on the relationships between different features—have evolved.

Overall, our results add to the body of knowledge showing that a small number of genomic traits—that includes utilization of the SD sequence mechanism—can be used to predict variation in minimum doubling times with surprising accuracy. Our findings demonstrate that measurements of SD sequence utilization outperform more commonly known associations such as the number of rRNA genes at this task. We believe that this is, in part, a consequence of the evolutionary inertia of different features [48]. In short, genome-wide usage of the SD sequence mechanism, like codon usage bias, requires hundreds of mutations to substantially alter and thus this trait will evolve much more slowly across a phylogeny when compared to more evolutionarily labile traits that rely on copy number variation such as rRNA and tRNA gene counts.

Like codon usage biases and in contrast to rRNA and tRNA gene counts, summary statistics based on SD sequence utilization do not require complete genome sequences and therefore may be *estimated* with partial genome fragments. The results and methods that we present here may thus have important applications in our understanding of novel, uncultivated genomes, environmental meta-genomic sequencing e orts, and the relationship between higher-order genome traits and growth strategies [50].

## Materials and Methods

### Data assembly

We first assembled a database of prokaryotic genomes from NCBI using the GBProks software (https://github.com/hyattpd/gbproks), including only ‘complete’ genomes in our download and subsequent analysis (accessed on: March 10, 2016). From the annotated GenBank files, we excluded pseudo-genes and plasmid based sequences from all subsequent analyses and proceeded to compile a data table with several traits for each genome. In addition to SD sequence utilization summary statistics described below, we applied RNAmmer to each genome in order to compile a list of ribosomal-RNA genes, and tRNAscan-SE to assemble a list of the tRNA genes [51, 52].

We wrote custom scripts to calculate the fraction of annotated coding sequences that begin with ‘ATG’, as well as the metric of codon usage bias (Δ*ENC* as described in [47]). For this latter metric, we first parsed the gene annotations to find ribosomal protein coding genes. We next computed the relative differences in codon usage bias between ribosomal protein coding genes and the rest of the genome, whereby:

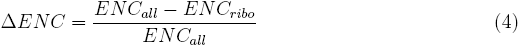

where ‘all’ and ‘ribo’ refer to all protein coding genes and ribosomal protein coding genes respectively. We altered the method used to calculate the ‘effective number of codons’ or ‘ENC’ from the one originally used by Vieira-Silva *et al.* (2010) to better control for GC content differences according to recent metric developed in our lab (manuscript submitted). The interpretation is the same, with values close to one occurring when ribosomal protein coding genes are very distinct in their codon usage bias patterns from the rest of the genome. By contrast, values close to zero occur when there is little codon usage bias separation between the genome and ribosomal protein coding genes.

For data on minimum doubling time, we downloaded the data table from Vieira-Silva *et al.* (2010), and paired each bacterial species with a complete genome from our database resulting in 187 matched species. To control for shared ancestry in subsequent analyses, we constructed a phylogenetic tree based off the rRNA sequences for this set of species. We first used RNAmmer to extract a randomly chosen 16S and 23S rRNA sequence from each genome, followed by MUSCLE (v3.8.31) on each individual rRNA to produce a multiple-sequence alignments [53]. These were concatenated together and we conducted a partitioned analysis using RAxML to construct a final tree. We performed 100 rapid Bootstrap searches, 20 ML searches and selected the best ML tree for subsequent analysis [54].

For the larger data-set, we instead relied on a previously computed high-quality phylogenetic tree published by Hug *et al.* (2016) [46]. We used custom scripts to match entries in this tree with genomes from our complete-genome database, and pruned away all species without a high-quality match resulting in 613 bacterial species in our final dataset that were used for subsequent analyses. For temperature annotations, we matched this set of 613 species to the ProTraits database using custom scripts, and restricted our analysis to species with temperature annotations exceeding a precision of 0.9 (equivalent to a FDR < 0.1) [55].

### Calculating summary statistics of SD sequence utilization

The calculation of Δ*I* is illustrated mathematically in the main text. Here, we only add that the calculation of the randomized sequences for all SD summary statistics is performed by first shuffling the upstream region of each gene between the region −30 to 0 (the first base of the start codon). Having shuffled each gene in this manner, we then performed the analysis as discussed in the main text for this shuffled ‘genome’ and repeat this calculation 500 times in order to derive null expectation for *f*_*SD*_, *f*_*aSD*<−4.5_ and *I*.

Next, we elaborate on our calculation of the other two methods for calculating SD sequence utilization. For each genome, we extract the −20 to −4 region upstream of the start codon for each gene. For *f*_*SD*_, we consider a gene as being SD-led if, in this defined region, any of the following motifs appear: ‘GGAA’, ‘GGAG’, ‘GAGG’, ‘AGGA’, or ‘AAGG’. We repeat this same process for 500 randomized ‘genomes’ where a randomized genome is defined as noted above (with the nucleotide region from −30 to 0 for each gene shuffled on a per-gene basis) prior to motif search.

For *f*_*aSD*<−4.5_, we perform a nearly identical procedure to the one listed above with the major difference being that instead of searching the upstream region of genes for particular motifs, we evaluate the hybridization energy between each 8 nucleotide segment contained within the −20 to −4 region and the putative aSD sequence defined as 5′-ACCUCCUU-3′ using the the ‘cofold’ method of the ViennaRNA software package with default parameters. If any sequence binds at a threshold of −4.5 kcal/mol or stronger (i.e. more negative Δ*G* values), we consider this gene to be SD-led.

### Phylogenetically generalized least squares

Throughout this manuscript, we utilize Phylogenetically Generalized Least Squares regression in order to mitigate the effects that arise from shared ancestry in statistical analyses. Our Phylogenetically Generalized Least Squares analysis relies on the most common null model, which assumes a Brownian motion model of trait evolution. For all statistical analyses presented in the paper, we use the R package ‘caper’ and perform a simultaneous maximum-likelihood estimate of Pagel’s λ, a branch length transformation, alongside the coefficients for independent variables of interest. All *p*-values that we report come from the F-test according to these results. For temperature analysis, we assigned ‘mesophiles’ and ‘thermophiles’ a value of 0 and 1 respectively and performed the equivalent fixed-effect analysis with Δ*I* as the dependent variable.

### Data availability and computer code

Data is provided as a supplementary file and all custom scripts and code that is sufficient to perform the analysis can be found at http://github.com/xxxx

## Acknowledgements

The authors wish to thank Thomas Stoeger for helpful discussions and critical reading of the manuscript, and Helio Tejedor for general computational support.

## Author contributions

AJH, MCJ and LANA conceived and designed the study. AJH collected the data and performed analysis. AJS contributed important preliminary results. AJH, MCJ, and LANA provided interpretation, and wrote the manuscript.

## Materials and correspondence

Correspondence and materials request should be addressed to

## Supplementary Information: Diversity of translation initiation mechanisms across bacterial species is driven by environmental conditions and growth demands

**Supplementary Table S1:** As in Table 1 of main text, instead showing results when Δ*f*_*aSD*<−4.5_ is used as the summary statistic for SD sequence utilization. Bold numbers in the far right column illustrate the variables with significant coefficients in the complete model (*p* < 0.001).

| Model for<br>min. doubling time | $R^2$ | Pagel's $\lambda$ | $\vert \Delta R^2\vert $ |
| --- | --- | --- | --- |
| Full model | 0.27*** | 0.94<br>[0.86,0.97] | - |
| $\Delta ENC$ | 0.12*** | 0.97<br>[0.93,0.99] | **0.13** |
| $\Delta f_{aSD < -4.5}$ | 0.09*** | 0.97<br>[0.93,0.99] | **0.06** |
| 16S gene counts | 0.06** | 0.98<br>[0.95,0.99] | 0.01 |
| tRNA gene counts | 0.06*** | 0.98<br>[0.95,0.99] | 0.02 |
| ATG start % | 0.02 | 0.98<br>[0.95,0.99] | < 0.01 |

**Supplementary Figure S1:**
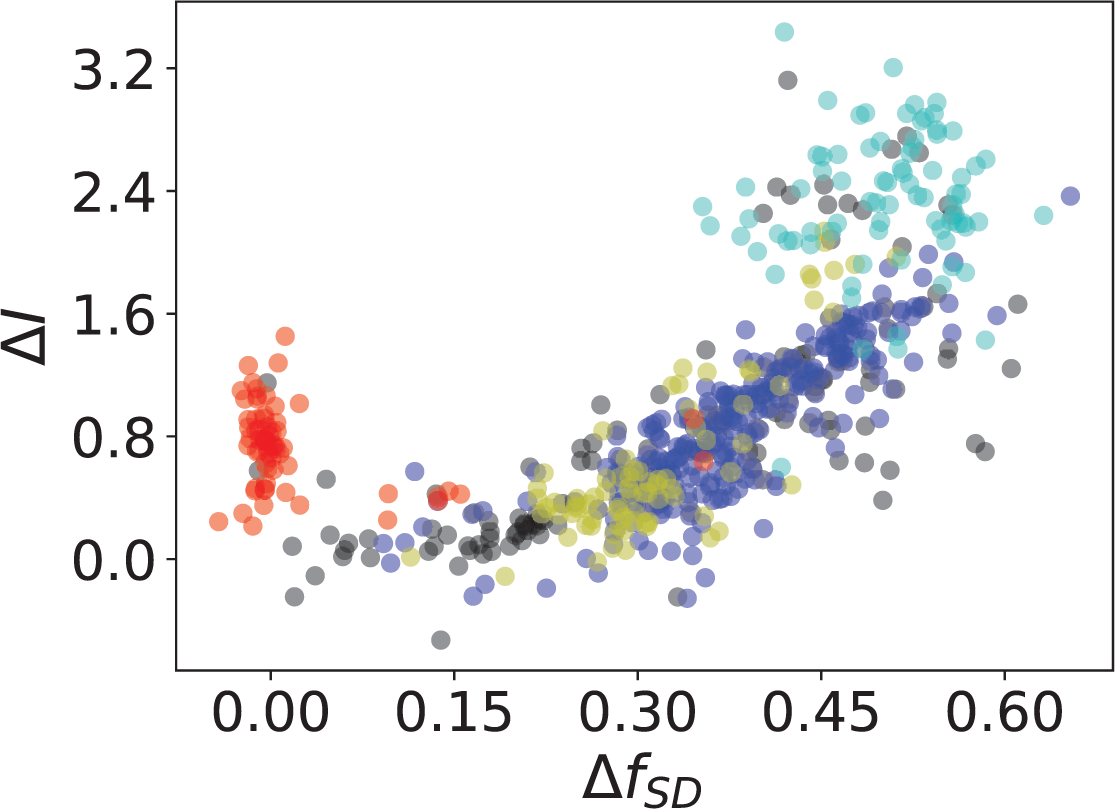
As in Fig. 2A (right) of main text, instead showing the relationship between Δ*f*_*SD*_ and Δ*I*.

**Supplementary Figure S2:**
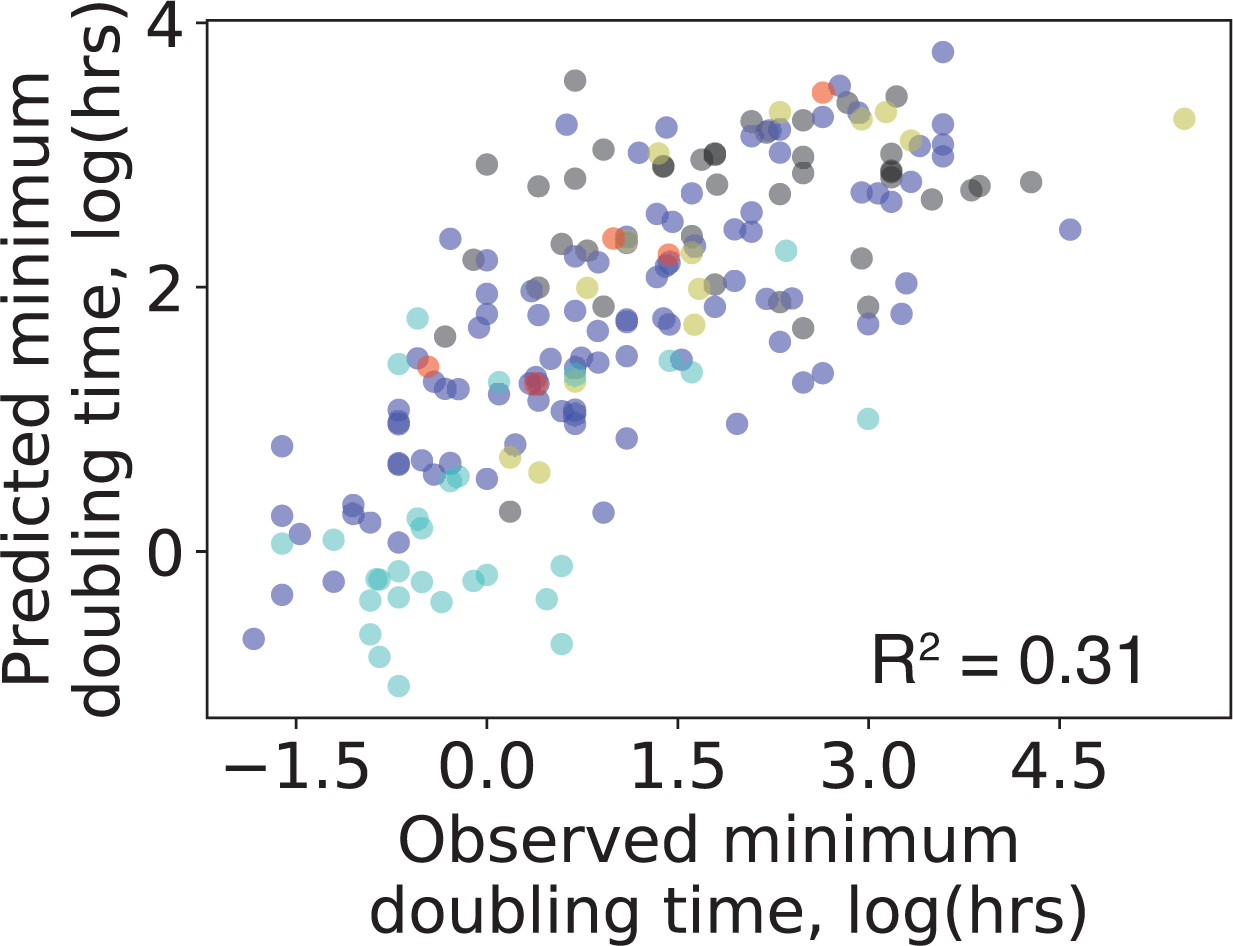
We show the observed and predicted values from a Phylogenetic Generalized Least Squares regression model using all predictors in Table 1. Species data points are colored according to phyla as in Fig. 2A.

**Supplementary Figure S3:**
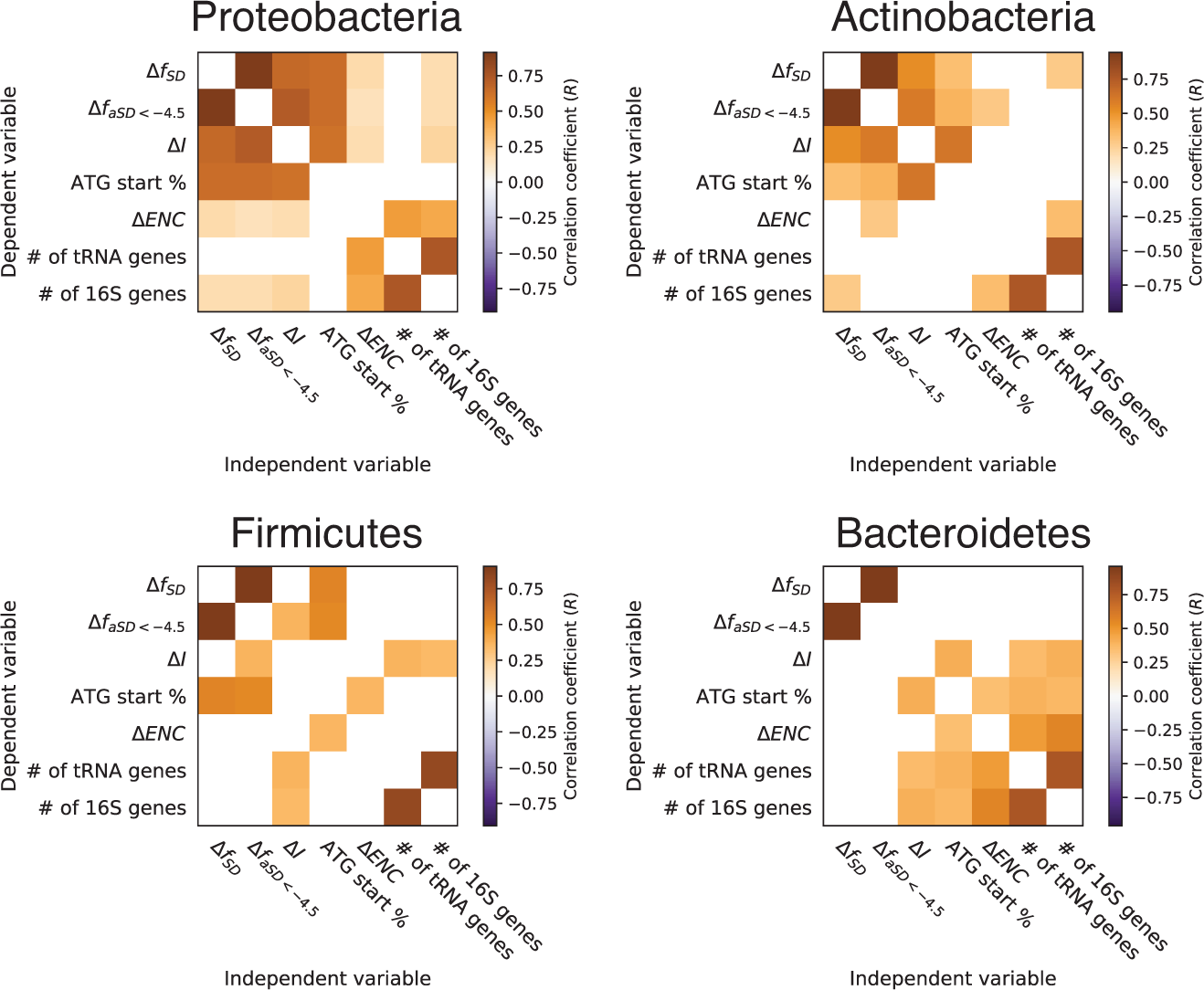
Correlation matrices as in Fig. 3A. We re-ran the analysis independently for each of the 4 major clades to illustrate the robustness of the conclusions to different groupings of species. Δ*f*_*aSD*<−4.5_ and Δ*f*_*SD*_ fail to show significant results for the Bacteroidetes phylum, by contrast Δ*I* uncovers these relationships.

